# A particle-filter framework for robust cryoEM 3D reconstruction

**DOI:** 10.1101/329169

**Authors:** Mingxu Hu, Hongkun Yu, Kai Gu, Kunpeng Wang, Siyuan Ren, Bing Li, Lin Gan, Shizhen Xu, Guangwen Yang, Yuan Shen, Xueming Li

## Abstract

Electron cryo-microscopy (cryoEM) is now a powerful tool in determining atomic structures of biological macromolecules under nearly natural conditions. The major task of single-particle cryoEM is to estimate a set of parameters for each input particle image to reconstruct the three-dimensional structure of the macromolecules. As future large-scale applications require increasingly higher resolution and automation, robust high-dimensional parameter estimation algorithms need to be developed in the presence of various image qualities. In this paper, we introduced a particle-filter algorithm for cryoEM, which was a sequential Monte Carlo method for robust and fast high-dimensional parameter estimation. The cryoEM parameter estimation problem was described by a probability density function of the estimated parameters. The particle filter uses a set of random and weighted support points to represent such a probability density function. The statistical properties of the support points not only enhance the parameter estimation with self-adaptive accuracy but also provide the belief of estimated parameters, which is essential for the reconstruction phase. The implementation of these features showed strong tolerance to bad particles and enabled robust defocus refinement, demonstrated by the remarkable resolution improvement at the atomic level.

## Introduction

Electron cryo-microscopy(cryoEM)is inaugurating a new era for structural biology by its capabilities to reveal atomic-resolution structures of biological samples under nearly natural conditions^1-3^. The cryoEM technologies are getting great attentions from both biologists and pharmaceutical companies, and leading to the construction of many cryoEM facilities in recent years. The recent achievements of cryoEM benefit from a series of technical breakthroughs since 2013^4-6^. The efforts on direct electron detector technologies, especially the electron counting, enabled efficient recording of atomic-resolution signals^7-10^. In the meantime, the development of computing algorithms, from motion correction to three-dimensional (3D) reconstruction and classification, are getting rapid progresses to obtain more accurate structural information hidden in strong noises of low-dose cryoEM images than ever^8,11-16^.

The increasing demand for general applicability and high throughput at atomic-resolution level is boosting new challenges for current cryoEM computing technologies. The general applicability often means high accuracy and robustness for datasets with various qualities. The application of a Bayesian-likelihood approach has shown advantages for the applicability, which introduced a statistical model to guide the iterative alignment processing in 3D reconstruction^14,17-19^. In each alignment step the orientation parameters including rotation and translation were estimated for the best matches evaluated by the likelihood of the experimental image against one or multiple given 3D references. The global goal of 3D reconstruction is to maximize the overall likelihood according to the Bayes rule. This statistical model makes the cryoEM 3D reconstruction more robust and automated than traditional methods. However, the high-dimensional parameter estimation is still a tricky but key factor in obtaining reliable results. The failure in reconstruction or classification, as well as issues like low resolution and over refinement, is usually caused by a bad parameter estimation, which has been recognized as an issue in traditional approaches. Moreover, the parameter estimation occupies a large portion of computing time in the entire processing, making it be a major target of computing acceleration. For some cases, extending the parameter estimation to higher-dimensional settings is necessary to obtain atomic-resolution reconstruction. However, it remains a challenge for both algorithm development and computing acceleration.

Owing to the importance of parameter estimation, many algorithms have been developed for cryoEM. The gridding and gradient algorithms, and a mixture of them, are popular in many published software. The gridding method is usually robust but with limited accuracy and high computing cost. Optimized versions of the gridding algorithm were thus developed, for example, an adaptive gridding algorithm with combination of coarse and fine grids used in RELION^14^. The gradient algorithms are well established in many fields to obtain fast computing performance. However, the robustness in global optimization and high-dimensional parameter estimation is always concerned. Therefore, the gradient algorithms were usually used for fast local parameter estimation, such as the conjunction gradient algorithm in FREALIGN^20^. Other more complicated algorithms are being considered. Recently, cryoSPARC implemented a stochastic gradient descent algorithm to obtain fast low-resolution model initialization, and combined a branch-and-bound algorithm to get rapid parameter estimation for high resolution^15^. However, current algorithms mostly focus on finding the best parameters, and lack of per-parameter evaluation of the estimation error. These also influence the robustness of high-dimensional parameter estimation.

Here, we implemented a particle-filter algorithm for single-particle cryoEM, which used statistical inference methods to find the complete solution of cryoEM parameter estimation. The particle-filter algorithm has been used to obtain optimal Bayesian estimation for nonlinear/non-Gaussian tracking problems^21-23^. It is a sequential Monte Carlo method, and uses random support points with weights to represent probability density function (PDF) of the parameter estimations. For cryoEM, the parameter estimation in the 3D alignment step can be described by a PDF that represents the belief of the estimated parameters. We designed a specialized particle-filter algorithm which estimated this PDF based on the likelihood function (LF) by a series of support points, and implemented it in the Bayesian approach^14,18^ for cryoEM 3D reconstruction. In the following, we applied the particle filter in four subspaces of the rotation, translation, defocus and structural state (3D classification). Then, we optimized the particle-filter algorithm and developed two weighting algorithms to improve the robustness of the 3D reconstruction. The current implementation highlights the self-adaptive adjustment for estimation accuracy, the tolerance to bad particles and the defocus refinement. The performance of the algorithm was demonstrated by the remarkable improvement in resolution of 3D reconstructions of several testing samples. These features make the particle-filter algorithm to serve as a novel framework of parameter estimation for future automation and large-scale application of cryoEM.

## Results

### A particle-filter algorithm for cryoEM parameter estimation

The alignment step in cryoEM 3D reconstruction is to estimate a set of high-dimensional parameters including orientations for each particle image. The measurement for the orientation parameters (the rotation and translation) has been formulated as the likelihood of an experimental image against a 3D reference of the target macromolecule^14^. Considering a high-dimensional parameter space consisting of three Euler rotation angles, two in-plane x-y translations, as well as some other parameters to describe the sample heterogeneity and imaging conditions, the measurements through this parameter space constitute a high-dimensional LF. The parameter estimation procedure, such as the classic gridding and gradient algorithms, can be modeled to search for the maximum likelihood, given by noisy and partial observations. Moreover, the LF contains more information which has not been fully exploited in classic cryoEM algorithms. The distribution of the likelihood, also known as the PDF of the parameter estimation, can be used to indicate the belief of the estimated parameters. The statistical information from the PDF provides a complete solution to the estimation problem, such as the measure of the accuracy at the single molecule particle level, and helps to achieve robust estimation in the high-dimensional space.

We implemented a specialized particle-filter algorithm to estimate the PDF used for cryoEM parameter estimation (see Method). The particle-filter algorithm performs estimation on the LF (Fig. 1a) by a set of random support points with associated weights (Fig. 1b). The distribution of the support points represents the PDF of the parameter estimation. In the beginning, a given number of the random support points are uniformly distributed in the entire parameter space for a good coverage to the whole LF for global search, called scanning phase (gray points in Fig. 1b). Then, a series of iterative phases are performed to gradually concentrate the random support points to the global optimum (colored points in Fig. 1b). In each phase, those sampling points located at higher likelihood regions will be assigned with higher weights and rapidly reallocated to more support points in this region (referred to as resampling process, see Method and Appendix I) in the following iterative phases. After obtaining rough estimations for the parameters in the global search, the local search is performed. The similar procedure is carried out as the global search but in a smaller region rather than the entire LF. This particle-filter algorithm can be used on any complicated high-dimensional LF with excellent robustness. We performed *ab initio* 3D reconstruction with an ellipsoid as the initial model for a dataset of the cyclic-nucleotide-gated (CNG) channel^24^ to test the algorithm, and the reconstruction correctly converged (the density maps in Fig. 1a). Meantime, the robustness of the particle-filter algorithm also provided the capability to optimize the defocus parameters together with the orientation parameters, which revealed the Z-height distribution of particles in the ice (discussed later).

**Figure 1.**
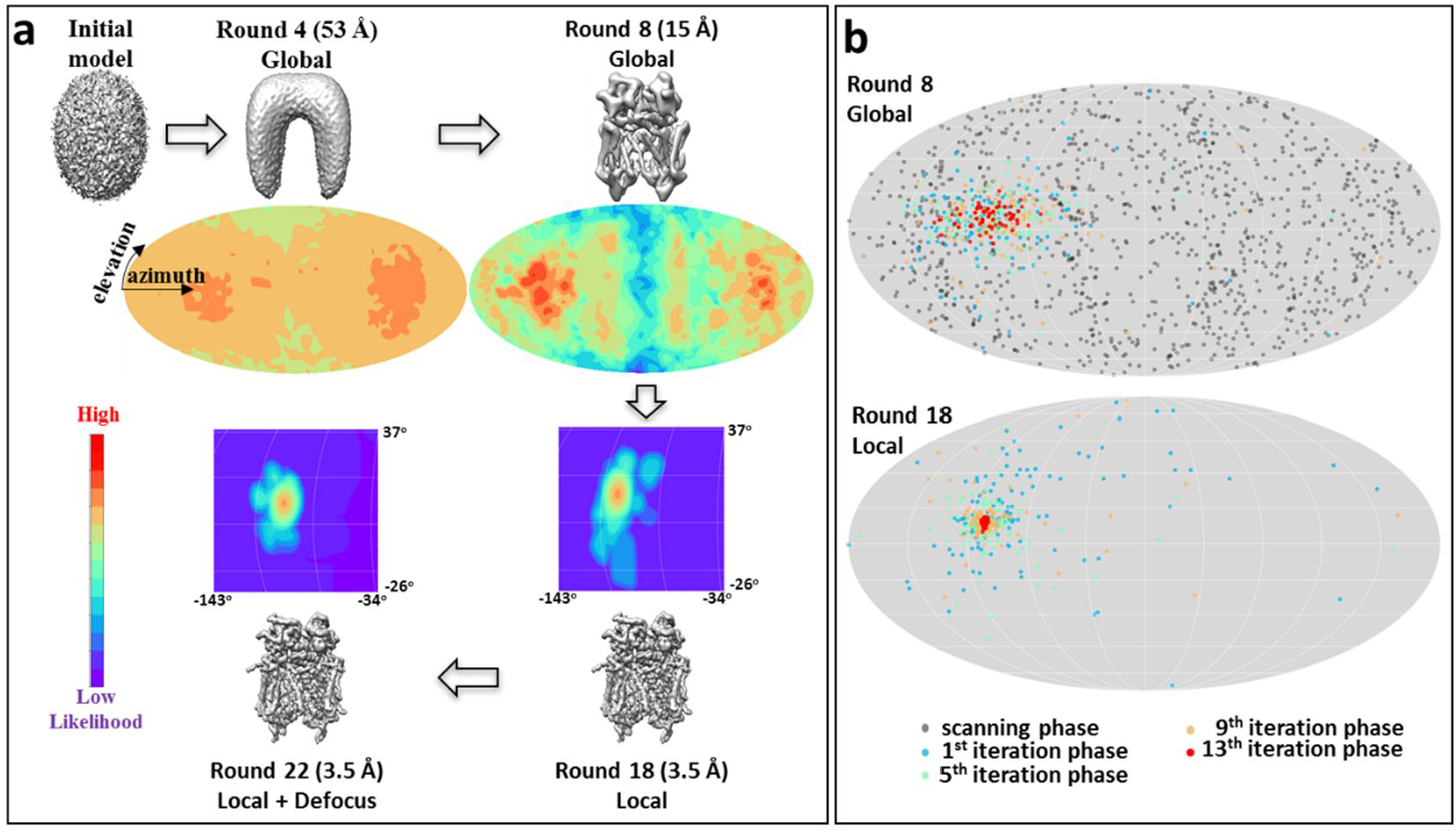
Illustration of support points in particle filter and the likelihood functions in rotation subspace of the CNG dataset. **a**) The 3D reconstructions and corresponding likelihood functions (the colored maps) in different rounds of 3D reconstruction. For the demonstration purpose, the 2D maps of the LFs are drawn by projecting the 3D LFs in 3D rotation subspace to the elevation-azimuth plane. Round 4 and 8 are global search performed at 53 Å and 15 Å resolution, respectively, and round 18 and 22 are local search performed at 3.5 Å resolution. In round 22, the defocus refinement was turned on. **b**) The support points used in the rotation subspace of round 8 and round 18 of **a**. Colors were used to indicate different iterative phases.

The structural state was considered as a classification parameter, corresponding to multiple 3D structures (see Method). Different from the rotation, translation and defocus, the structural-state parameter is discrete in our current algorithm, and the number of support points is set to be equal to the number of classes. We tested the 3D classification of the CNG datasets using the ellipsoid above as the initial model, and observed the classes losing part of the densities in the trans-membrane region of the CNG channel (Suppl. Fig. 1a), which agreed with our previous observation. This result confirmed the robustness of the particle filter in 3D classification from a simple initial model.

The initialization and the resampling of the particle filter needs sampling points with uniform or a certain distribution of the rotation parameters. This is a very difficult task using the Euler-angle system. Alternatively, we used the unit quaternion to describe the rotation operations in the implementation of the particle filter (see Method).

### Optimize the number of support points for fast and robust parameter estimation

The particle-filter algorithm uses a set of random support points to perform iterative updates based on the LF. As the number of total support points is directly proportional to the computing cost, it is necessary to optimize the number of support points to obtain fast computing performance. However, the selection of the number of supporting points depends on the complexity of the LF. Too few points will reduce the coverage to the LF and may lead to the local optimum. Optimizing the number of support points needs to balance the robustness and the computing performance.

In current implementation, the particle filter was applied to four subspaces, including the rotation, x-y translation, defocus, and structural state. The total number of support points is the product of those in all subspaces. Except for the structural-state subspace whose number of support points is equal to the number of classes, we evaluated the influences of different numbers of support points in other three subspaces. The Fourier Shell Correlation (FSC) curves were used as a criterion. If the parameter estimations on most particles go wrong or fall into the local optimum owing to too few support points, the resolution or the FSC curve of a 3D reconstruction is expected to become worse.

The LF in the rotation subspace has the most complicated form with many peaks (Fig. 1a), so that a large number of support points are put on the rotation subspace. In global search, our tests showed that ∼10,000 support points can give a good coverage to the rotation subspace in the scanning phase (Fig. 1b). If considering the symmetry of the molecule particles, the number of support points can be reduced by the number of symmetric operations. After the first iterative phase, the support points usually will be rapidly concentrated to the regions near the global optimum. Hence, ∼100 points are enough in the following iterative phases (Suppl. Fig. 2a). In local search, the estimation is always limited in a small region, and ∼100 points are empirically enough for the rotation subspace (Fig. 2a and Suppl. Fig. 2a). The LF in the translation and defocus subspace has a simple single-peak shape (Fig. 2b and c), and hence the number of support points can be much less than that for the rotation. For a translation estimation within 25-pixel radius, ∼70 support points are typically enough in the scanning phase. And then with the rapidly converged points, the number can be further reduced to ∼9 (Fig. 2b and Suppl. Fig. 2b). For defocus, rough values with astigmatism should already be available, such as that from CTFFind3^25^. In current implementation, only local defocus search without considering astigmatism is performed. ∼9 support points can give the correct result (Fig. 2c). Therefore, except for the scanning phase in global search, later phases of the parameter estimation require only a small number of support points for providing a correct estimation, which are typically ∼125, ∼9 and ∼9 points for the rotation, x-y translations and defocus dimension, respectively.

**Figure 2.**
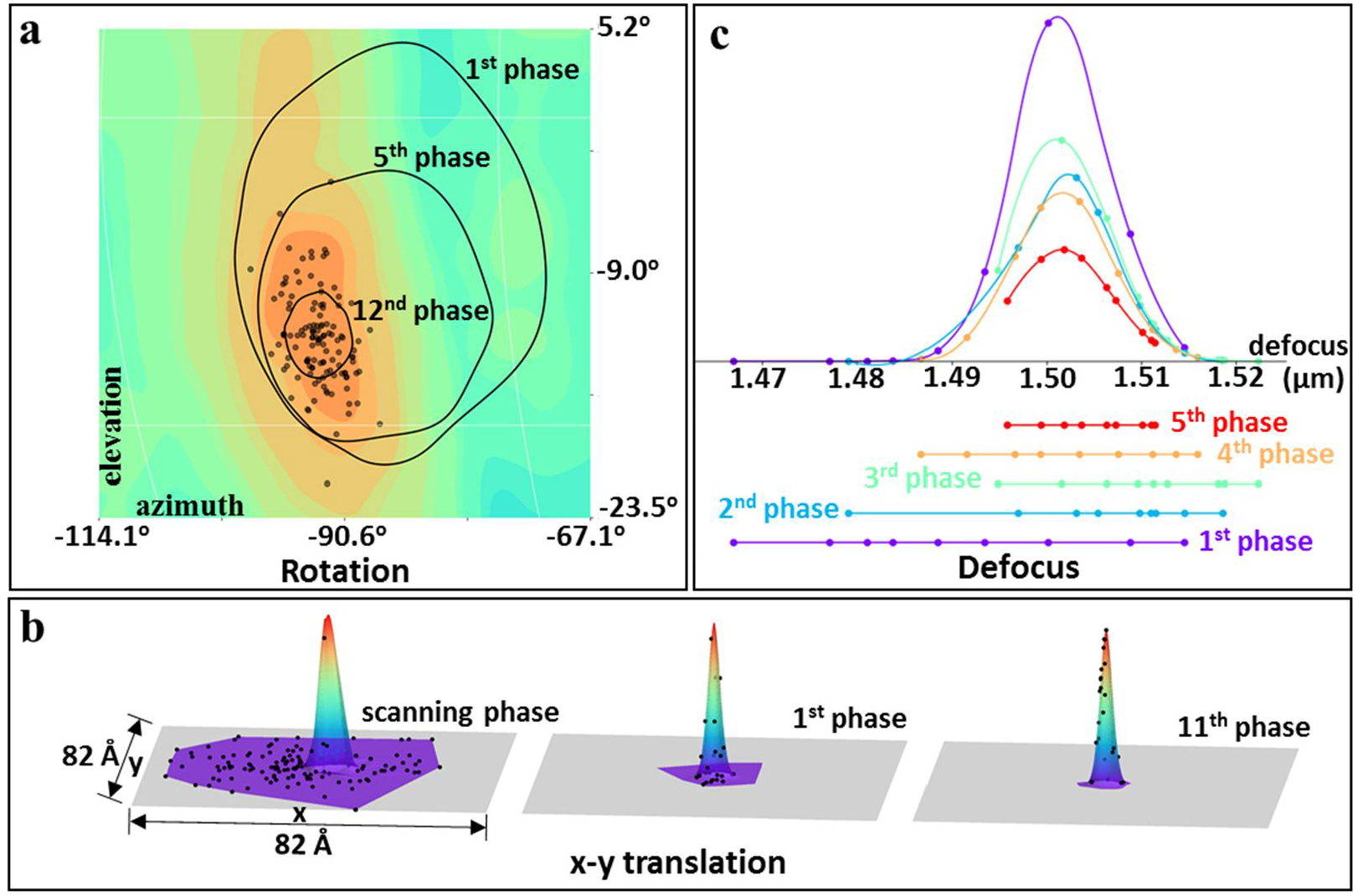
Supports points in three subspaces. **a**) the rotation subspace. The support points in rotation subspace were plotted in the elevation-azimuth plane by projecting all support points in 3D space into this 2D plane. The map of local LF (drawn in the same way as that for Fig. 1a) are shown in the background. The smallest cycle indicates the range of 1x standard deviation of the plotted support points. The other two cycles indicates 1x standard deviation of point distribution in the earlier phases. **b**) the x-y translation subspace. The gray plane is the x-y plane. The peaks are the fitting of weights of the support points. The support points in the scanning phase and later phases are shown. **c**) the defocus subspace. The black horizontal line is the defocus axis. The colored curves are the fitting of the weights of the support points. The points on the colored horizontal lines show the defocus distribution of the support points. Colors were used to specify different iterative phases.

### Self-adaptive accuracy for parameter estimation

The parameter estimation accuracy is important for the obtainable resolution in 3D reconstruction, which varies with the image quality in terms of the signal-to-noise ratio. In the grid algorithm, a finer grid always yields more accurate estimation, but tends to increase unnecessary computations when the reference model is in low resolution. Therefore, some prior knowledge is needed to achieve a balance between parameter accuracy and computing cost. The particle filter shows advantage in such problems by self-adaptive adjustment for parameter estimation accuracy.

If an estimation of a parameter has a large confidence, the local LF around this parameter exhibits as a high and sharp peak. The support points on the peak will obtain high weights (proportional to the likelihood, see Method), and concentrate dense points to its adjacent region (Fig. 2 and Suppl. Fig. 3a) by the resampling procedure. The dense support points provide fine coverage for this peak area to achieve accurate estimation. Otherwise, the LF is flat, which often happens at the low-resolution stage of the first several rounds of 3D reconstruction (Fig. 1a), or when the particle image does not match well with the 3D reference. Consequently, the support points will be distributed in a large area and even on multiple peaks (Suppl. Fig. 3b), which provides a coverage to a large region but with low sampling accuracy. This brings advantages of searching more regions to increase the possibility of finding the global optimum.

**Figure 3.**
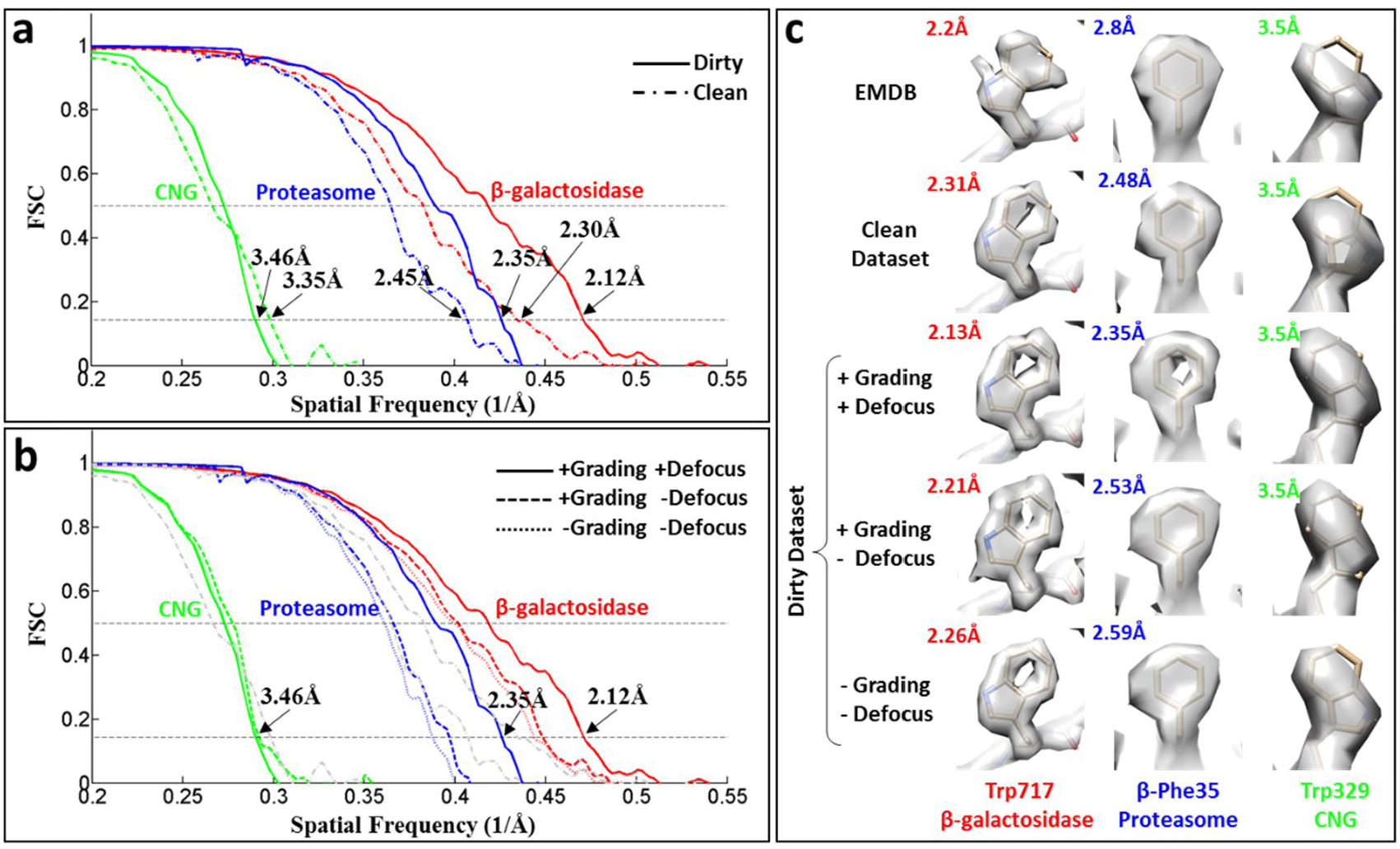
Resolution comparisons among different processing options for three datasets. **a**) FSC curves of the dirty and clean datasets. **b**) FSC curves with various options of the particle grading and defocus refinement. The grey curves are the FSC curves of the clean dataset shown in **a**. **c**) Selected side chains with different processing options. See **Suppl. Fig. 4** and **5** for more comparisons.

When the particle filter uses a fixed number of support points, the computation cost can be kept constant. In such cases, higher estimation accuracy can be achieved by optimizing the distribution of the random support points.

### Improve tolerance to bad particles by particle grading and distribution weighting

Identifying and removing bad particles to improve the resolution of 3D reconstruction is a difficult and time-consuming task. The major problem is that it is difficult to define the “bad” particles in a quantitative way. A possible characteristic of a bad particle is the lower “stability” during parameter estimation than that of the good particles, whose parameters are sensitive to any changes of noises or 3D references. This characteristic has been used to find the bad particles, such as a random-phase 3D classification method^26^ to count the frequency of a particle jumping between different 3D classes. However, so far there isn’t a simple and quantitative way to perform such estimation. As discussed in the previous section about the self-adaptive accuracy, the distribution of support points varied with the confidence of the parameter estimation. The low stability may be described by the low confidence here. Therefore, estimating the distribution of support points provides a statistical way to quantify the estimation confidence. Accordingly, we implemented two methods to utilize this information to adjust the contribution of each particle to the reconstruction (see Method). First, the reciprocal of the standard deviation of the support point distribution was used as a weighting factor for each particle image used in the reconstruction, named particle grading. Second, each particle image was inserted into the reconstruction for *N* times (empirically, *N* = 100 in our current implementation) with parameters following the distribution obtained from support points, named distribution weighting. If the distribution is wide, the contribution of the particle image is equivalent to be diluted, vice versa. Therefore, the second method is an inexplicit weighting algorithm.

To test the weighting algorithms, we used three high-resolution datasets, the proteasome^27^ (Entry code: 10025) and β-galactosidase^28^ (Entry code: 10061) from EMPIAR^29^, and the CNG dataset from our previous work^24^. In the published results, the initial datasets were subjected to a series of screening processes including 2D or 3D classifications to remove “bad” particles. And only less than 50% particles were selected and used in the final reconstruction. This ratio might be much lower, even 5∼10%, in some cases. Now our particle grading and distribution weighting can automatically deal with the “bad” particles so that no additional efforts are needed to filter bad particles. After the particle picking, we only performed a simple 2D classification to remove some wrong particles, ∼10% of the initial particles, which were mostly ice contaminations picked by automated particle picking. Then, 211,826 CNG particles, 112,412 proteasome particles and 89,857 β-galactosidase particles, simply called dirty datasets, were directly subjected to 3D refinement using the particle-filter algorithm. As a control, we also calculated the 3D reconstructions from the particles selected in the published works, simply called clean datasets, 87,149 CNG particles, 49,954 proteasome particles and 38,965 β-galactosidase particles.

As expected, the FSC curves of two dirty datasets (solid lines) of proteasome (blue curve) and β-galactosidase (red curves) show much better resolutions than the corresponding clean ones (dashdotted lines) (Fig. 3a). The significantly improvement of the map quality confirmed the resolutions (the second and third rows of Fig. 3c). Some aromatic rings in β-galactosidase show clearer holes in the center than both the clean dataset and the published map (EMDB entry code: 2984) (Suppl. Fig. 4). Especially, the Phe35 in β subunit of the proteasome showed a hole in its aromatic ring (Fig. 3c), which demonstrated the resolution improvement of almost half angstrom from 2.8 Å of the EMDB map (EMDB entry code: 6278). These results indicate that many good particles were removed together with the bad ones during the screening processes in the published works. By our particle filter algorithm, the bad particles were automatically set with low contribution to the final reconstructions, and hence more good particles can be used to improve the resolution. Therefore, the grading and distribution weighting improved the tolerance to bad particles. To further investigate the contribution from the particle grading itself, we performed 3D refinement with and without particle grading (defocus refinement was disabled for this test) for the dirty datasets. The FSC curves (Fig. 3b) with particle grading (dashed lines) showed improvements relative to those without grading (dotted lines), which demonstrated the contribution of the particle grading. The contribution of distribution weighting wasn’t compared, because it was part of the reconstruction and cannot be turned off.

**Figure 4.**
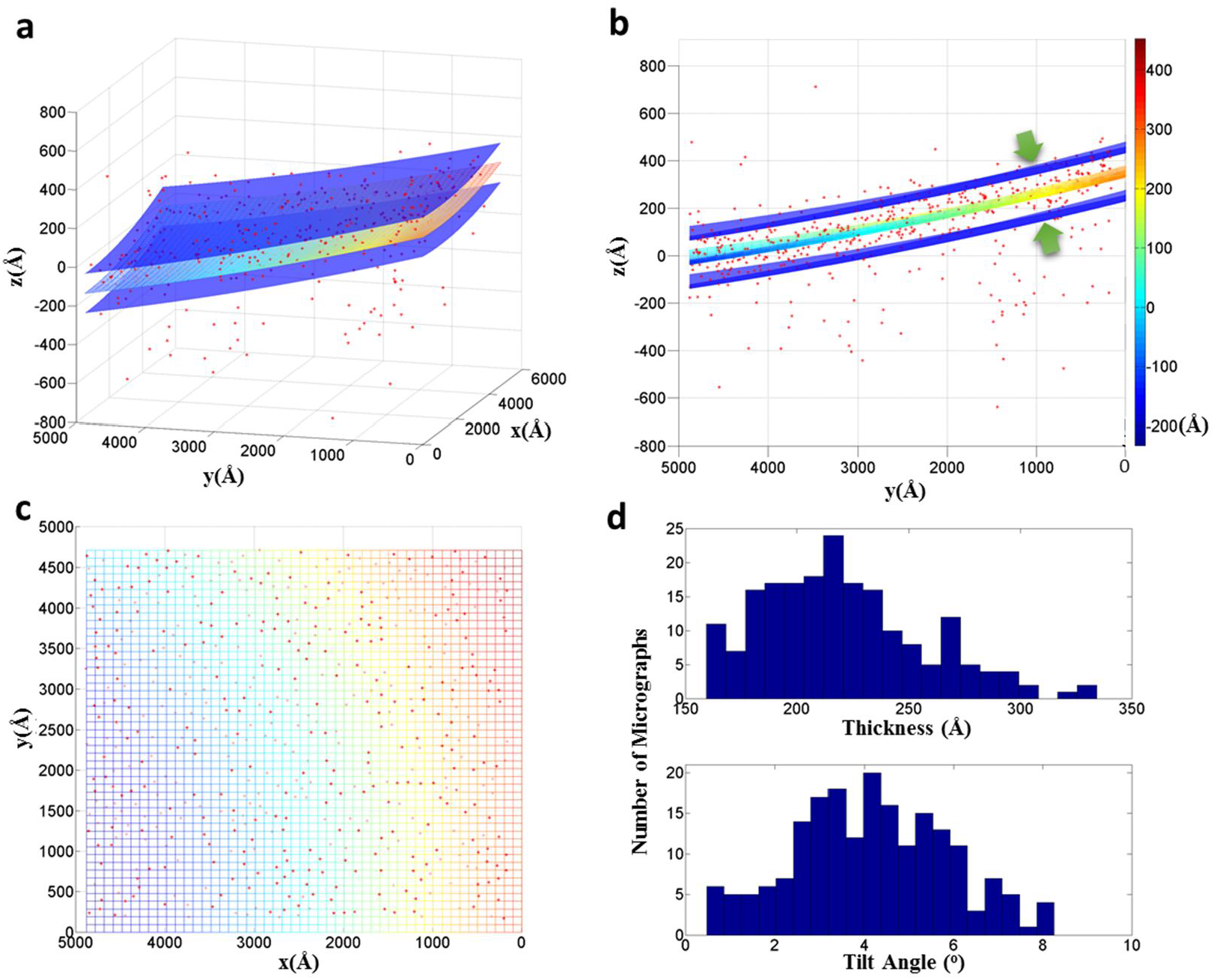
Z-height distribution of the proteasome particles in the ice. 3D plot of particle positions in one typical micrograph is shown in **a**) stereo view, **b**) side view along x axis and **c**) top view along z axis. A fitting to a quadric surface (the colored surface) showed that 75% particles located in a thin layer (between two blue surfaces) with 4.5° tilt angle and ∼200 Å thickness. **d**) histogram of the tilt angles and thicknesses measured by quadric surface fitting from all 196 micrographs in the proteasome dataset.

For the CNG dataset, we performed the same tests as those for the other two datasets, but no significant improvement was observed. On the other hand, the bad particles in the dirty dataset also didn’t cause degradation of the resolution, and we can even see some subtle improvements in the map of the dirty dataset (the third column of Fig. 3c). In our current and previous works, it was observed that some CNG complexes had a flexible subunit. Therefore, the flexibility of CNG complex might limit the obtainable resolution.

### Defocus refinement for accurate Z height measurement of macromolecules in ice layer

Protein particles may have different Z heights (assuming the incident beam is along Z axis) in the same ice layer of cryoEM grid, which results in varied defocus for particles in the same micrograph. The defocus variation can be as large as several hundreds of angstroms due to thick ice or stage tilt, and introduce significant phase error to high-frequency signals during the correction for contrast transfer function (CTF). Therefore, determining accurate defocus parameter is essential for obtaining atomic resolution better than 3 Å. Benefiting from the robustness and good computing performance of the proposed particle-filter algorithm, we are able to extend the parameter estimation to the defocus based on the signal from each single particle.

We tested defocus refinement using the three datasets of CNG channel, proteasome and β-galactosidase. Both the proteasome and β-galactosidase showed remarkable improvements in resolution indicated by FSC shifts of 0.1∼0.2 Å to 2 Å resolution (Fig. 3b). The density maps were also significantly improved, which was illustrated by the clearer holes in some aromatic rings (the third and fourth rows of Fig. 3c). This result confirmed that high-frequency signals are more sensitive to the phase error due to the inaccurate defocus value. For the CNG dataset, we didn’t see improvement of resolution with and without defocus refinement (green solid and green dash line, respectively, in Fig. 3b). A possible reason is that the reconstruction at the resolution around 3.5 Å is not sensitive to the phase error. And it is also possible that the ice of the CNG sample is thin and flat, and hence there is no large defocus variation.

Especially, the proteasome dataset shows the largest improvement from 2.53 Å to 2.35 Å with defocus refinement. To investigate the possible reason, we did a 3D plot for the particle positions of each proteasome particle, where the Z coordinates are the relative defocus variation from the defocus measured by the whole micrograph (Fig. 4a-c). We observed that most particles distributed in a thin layer of 160∼300 Å thickness (Fig. 5d). And most micrographs have a large tilt of 1° ∼8° (Fig. 4d). The defocus difference from a corner to another can be more than 400 Å for a 5° tilt. Therefore, such large sample tilt may explain the remarkable improvement after defocus refinement. Moreover, we found that a quadric surface gave a better fitting to the particle positions than a plane, which might indicate that the ice layer was slightly curved (Fig. 4c). Such a result also agrees with the observations of ice deformation under electron radiation^11,30^.

## Discussion

The current cryoEM technologies have made atomic resolution more and more obtainable than ever. However, the requirements for large-scale applications of higher resolution better than 3 Å are still challenging for future cryoEM technologies. Many bottlenecks, such as low-quality samples, dependence on user experiences and measurement errors in imaging parameters, still exist. Under this background, we implemented a specialized particle-filter algorithm to improve the robustness and accuracy of the single-particle 3D reconstruction. The particle filter, a statistical inference method, has demonstrated superior accuracy and robustness with affordable computations in many engineering applications^23^. The basic idea of particle filters is to use a series of support points with corresponding weights to represent complex probability distributions. In the case of cryoEM 3D reconstruction, we used the support points to represent the PDF of estimated parameters constructed on the LF in the high-dimensional parameter space. By optimizing the particle filter in the cryoEM parameter space, involving three subspaces of the rotation, xy-translation and defocus, we were able to achieve low-complexity but accurate parameter estimation for atomic-resolution 3D reconstruction. The application on 3D classification initialized from a simple ellipsoid model also proved the robustness of our particle-filter algorithm. The performance of this particle-filter algorithm was then demonstrated by significantly improved resolutions for two datasets of proteasome^27^ and β-galactosidase^28^ over the published results (Suppl. Fig. 5). Especially, we pushed the resolution of the proteasome dataset to ∼1.1x physical Nyquist frequency. This also demonstrated that our particle-filter implementation is able to reveal the super-resolution signal beyond the physical Nyquist frequency of K2 counting camera (Gatan Company) through accurate 3D parameter estimation.

The particle-filter algorithm provided an effective metric for the confidence of parameter estimates, which is quantified by the standard deviation of the distribution of the support points. This confidence is actually comparable among different particles, and hence can be used for identifying bad particles. This is a unique function benefiting from the statistical method used in particle filters. We utilized this information through two weighting algorithms, the particle grading and distribution weighting. The tests on the three datasets show excellent tolerance to bad particles, so that no extra processing, such as recursive 2D and 3D classification, is needed to remove bad particles. The 3D classification has been intensively used in a trial-and-error way to find and remove bad particles in many published works. Such classification processes strongly depend on user experiences and available computing resources, and often do not work well for the beginner. The particle grading and distribution weighting not only provide a solution for this kind of problems by reducing extra processing to push the resolution, but also enhance the robustness of the 3D reconstruction. Together with self-adaptive accuracy, the particle grading and distribution weighting also make the 3D reconstruction procedure more automated.

For the efforts to push the resolution to the atomic level, the defocus variation must be fully considered. In the resolution range better than 3 Å, the oscillation of the CTF becomes severe. Small errors in the defocus value can cause large phase errors in 3D reconstruction of high-frequency signals. Therefore, the defocus change caused by Z height variation of particles in thick or tilted ice must be precisely determined for each particle image. Programs, such as CTFTilt^25^, have been used to measure the sample tilt on the whole micrograph. A recently published software package GCTF enabled the determination for local defocus^31^. But all these software is based on the Thon ring signal from both particles and surrounding ice, which cannot handle the per-particle Z height change in the ice. The robustness of high-dimensional parameter estimation by particle filters enabled the defocus refinement together with other parameters, and our tests showed significant resolution improvement for the reconstruction in the resolution range better than 3 Å.

## Acknowledgements

This work was supported by funds from The National Key Research and Development Program (2016YFA0501102 and 2016YFA0501902 to X.L.), National Natural Science Foundation of China (31570730 to X.L., and 61672312 to G.Y.), Advanced Innovation Center for Structural Biology (to X. L. and Y. S.), Tsinghua-Peking Joint Center for Life Sciences (to X. L.) and One-Thousand Talent Program by the State Council of China (to X. L. and Y. S.). We acknowledge National Supercomputing Center in Wuxi and Tsinghua University Branch of China National Center for Protein Sciences Beijing for providing facility supports in computation.

## Contributions

X.L., Y.S. and G.Y. initialized the project, Y.S., M.H., X.L., H.Y. and K.G. designed the algorithm, M.H. and H.Y. implemented the algorithm, M.H. and H.Y. designed and wrote the major part of the program with full functions of 3D reconstruction, K.W., S.R., B.L., L.G. and S.X. wrote part of the program. M.H. and X.L performed the tests. X.L. wrote the manuscript, and all authors revised the manuscript.

## Competing financial interests

The authors declare no competing financial interests.

## Method

### Particle filter algorithm

For single-particle cryoEM 3D alignment, the core step is to determine a set of, possibly high-dimensional, parameters *x* = {*Φ*, *t*, *ζ*, *µ* … } for each particle image, where *Φ* stands for the rotation, *t* stands for the translation in the image plane, *ζ* stands for the defocus or the particle position along incident beam, *µ* stands for a underlying structural state, and other parameters in the cryoEM system can also be considered. In the present work, we will focus on determining the first four sets of parameters, i.e., rotation, translation, defocus and structural state for 3D classification. The corresponding method can also be extended to the determination of more parameters.

With the attempt to improve the robustness for high-dimensional parameter estimation, we expressed the belief of parameter *x* by its PDF, *p*(*x|𝒳*, *V*), where *x* stands for the experimental particle image, and *V* stands for the 3D reference. If the 3D classification is performed, *V* will contains a set of underlying reference structures. Such probabilistic description not only gives the estimation of *x*, but also has the advantage in obtaining the estimation error of *x* for each experimental image against the given 3D reference.

We next introduced a particle-filter algorithm^21-23^ to estimate *p*(*x|𝒳*, *V*). The particle filter is a Monte Carlo method representing the required posterior PDF by a set of random support points with associated weights. In our cryoEM implementation, the particle-filter algorithm aims to construct a PDF, *q*(*x|𝒳*, *V*), represented by a set of *N* random support points {*x*_*i*_}_*1≤i≤N*_ in the high-dimensional cryoEM parameter space, to approximate *p*(*x|𝒳*, *V*). The distribution *q*(*x|𝒳*, *V*) was usually called importance density in the engineering applications of the particle filter^21^.

A brief introduction for the implemented algorithm is as follows.

The particle filter undergoes a series of iterative phases to reach

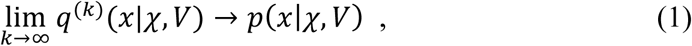

where *k* is the iterative number, and *q*^(*k*)^(*x|𝒳, V*) is the evaluated PDF from previous *k - 1* iterations. In the first iterative phase, *q*^(1)^(*x|𝒳, V*) is usually initialized as a uniform distribution. Then, *q (x|𝒳, V*) propagates as update from *q*^(*k*)^(*x|𝒳, V*) to *q*^(*k+1*)^(*x|𝒳, V*) to reach a precise estimation of *p*(*x|𝒳*, *V*).

In the beginning of *k*^th^iterative phase, we assume that *q*^(*k*)^(*x|𝒳, V*) is known and represent it by *N* support points 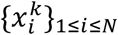 as

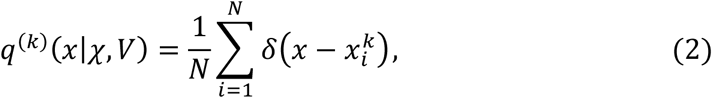

where *δ* is Dirac function and 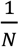 is the normalization factor. The difference between *q*^(*k*)^(*x|𝒳, V*) and *p*(*x|𝒳*, *V*) on the support points 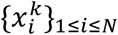 is formulated as

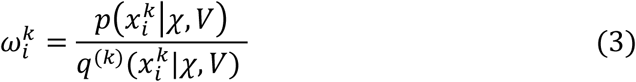

which is also referred to as associated weight of support point 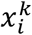. When *q*^(*k*)^(*x|𝒳, V*) tends to *p*(*x|𝒳*, *V*), 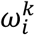 will tend to 1. Therefore, 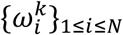 can guide us to generate a new set of supporting points 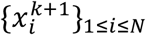 which represents a new distribution *q*^(*k*+1)^ (*x|𝒳, V*) with the better approximation to *p*(*x|𝒳*, *V*). To implement Eq. (3), we considered the Bayes’ theorem and got

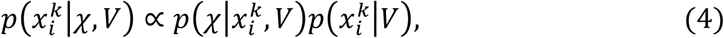

where 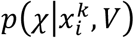 was the likelihood^14^ between experimental image *𝒳* and the projection of *V* with parameters 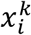 and we chose the prior to be

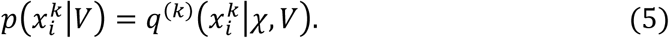

By substituting Eq. (4) and (5) into Eq. (3), the weight can be shown to be

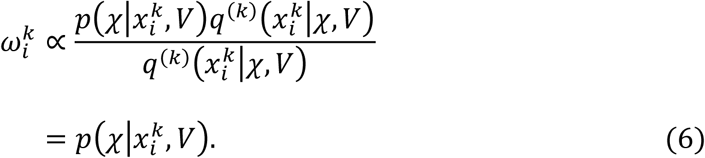

The normalized weight is used so that 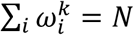 and henc

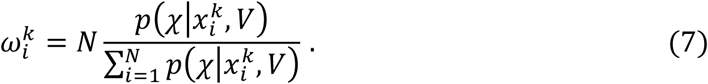

By now we obtain a set of support points associated with weights,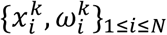 which approximates *p*(*x|𝒳*, *V*) in *k*^th^ iterative phase.

Then, we introduced a procedure called resampling to generate a new set of *N* support points 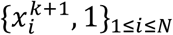 with equal weights from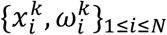. The basic idea is to concentrate the support points to the points with high weights. Therefore, if a support point in 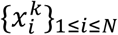 has a high weight, it will be divided into multiple resampled support points, while the points with small weights will be eliminated. In the end, the total number of support points is kept unchanged. A pseudo-code description of the resampling procedure is given in Appendix I. The newly generated support points 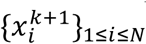 represent the updated distribution function in (k + 1)^th^ iterative phase as

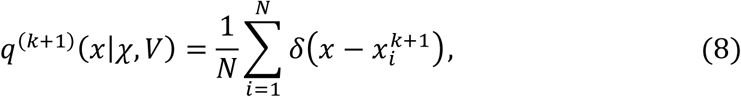

From Eq. (2) to (8), we developed a particle-filter algorithm specialized for cryoEM parameter estimation. A pseudo code is also provided in Appendix II to describe the detailed computing procedure of the particle-filter algorithm.

As a summary to the algorithm, the particle filter achieves the approximation to *p* (*x|𝒳, V*) by iteratively updating a discrete distribution represented by a set of support points. The updating procedure is guided by the associated weights of support points in each iterative phase. As described in Eq. (6), it is seen that these weights actually perform measurements on the likelihood function *f*(*x*) = *p*(*x|𝒳, V*) in the cryoEM parameter space.

The parameter space of *x* includes four subspaces in current implementation, i.e., rotation, translation, defocus and structural state, each of which has different physical meaning from others. Therefore, we performed particle filter in the four subspaces to estimate PDFs *p*(*Φ| 𝒳, V*), *p*(*t|𝒳, V*), *p*(*ζ|𝒳, V*) and *p*(*µ| 𝒳, V*) for the rotation, translation, defocus and structural state, respectively. Currently, *µ* is a discrete parameter indicating one of a set of structural states (3D classes). Then, the overall parameter estimation for *x* is approximated by the product of the marginalized PDFs

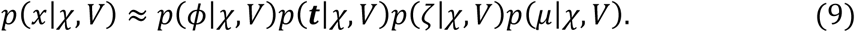

To achieve Eq. (9) by particle filter, we need to know the likelihood to calculate the weights of support points according to Eq. (6) and (7). In the following, we used the rotation parameter as an example to derive the likelihood function in the rotation subspace. Based on the total probability theorem, the likelihood in the rotation subspace can be calculated as follows

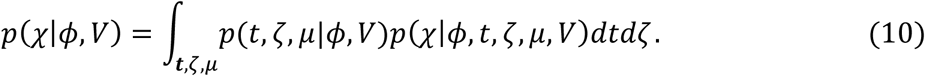

Assuming that the four subspaces of the rotation, translation, defocus and structural state are independent to each other, the middle items in Eq. (10) can be approximated as following

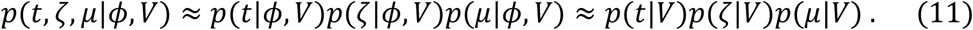

By choosing importance density *q*^(*k*)^(*x|*𝒳, *V*) to be the prior *p*(*x|V*) as Eq. (5), Eq. (11) can be estimated in the *k*^th^ iterative phase as

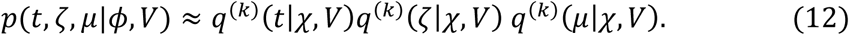

Substituting Eq. (2) and (12) into Eq. (10), and performing the same derivations for the other three parameters, we obtained the likelihood in the discrete form as

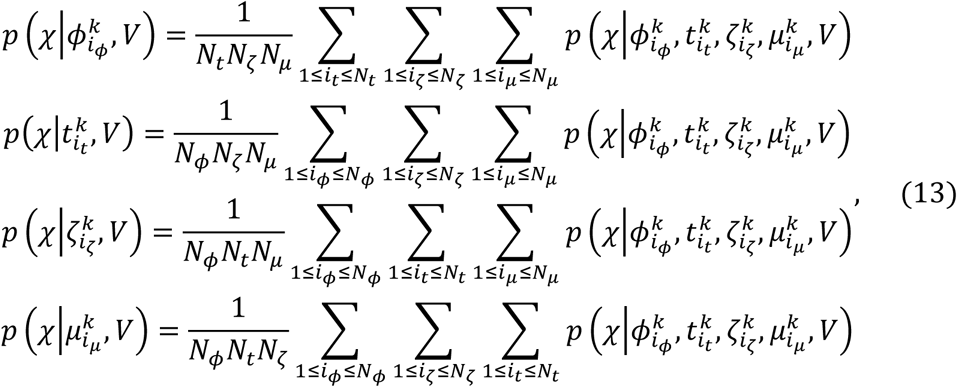

where *N*_*Φ*_, *N*_*t*_, *N*_ζ_ and *N*_*µ*_ are the number of support points 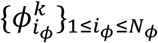, 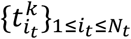, 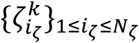 and 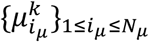 for rotation, translation and defocus, respectively.

### Particle grading and distribution weighting

For a dataset { 𝒳 _*l*_}_*1≤l≤L*_ with *L* particle images, the particle filter is carried out for each particle image 𝒳 _*l*_ to estimate the PDF *p*(*x|*𝒳_*l*_,V) represented by *N* support points {*x*_*il*_}_1≤*i*≤*N*_. Then, the 3D reconstruction is calculated as following

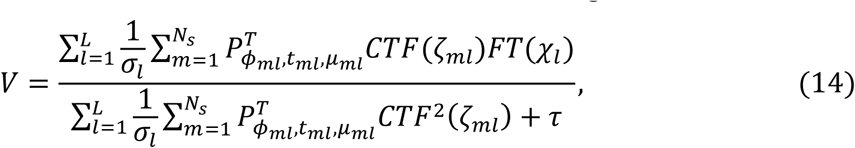

Where *σ*_*l*_ is the standard deviation of support points {*x*_*il*_}_1≤*i*≤*N*_, {*Φ*_*ml*_, *t*_*ml*_, *ζ*_*ml*_, *µ*_*ml*_} = *x*_*ml*_ ∈ {*x*_*il*_}_1≤*i*≤*N*_, *N*_s_ is the number of support points used in the reconstruction, *FT* is Fourier transform,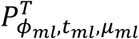 is an operation to place the Fourier transform of 𝒳 _*l*_ into the Fourier transform of the 3D reconstruction, *CTF*(*ζ*_*ml*_) is the contrast transfer function, and *τ* is a Wiener filter factor.

Here, we considered two weighting strategies in the 3D reconstruction described by Eq. (11), which is derived from *p*(*x|* 𝒳 _*l*_, *V*). First, the reciprocal of the standard deviation of support points,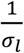, was used as an explicit weighting factor for image *ζ* _*l*_, called particle grading. A large *σ*l indicates low confidence for the parameter estimation. Consequently,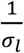 will reduce the contribution of the corresponding particle image xl to the final reconstruction, and *vice versa*. In current implementation, *a*_*l*_ is only calculated from the support points of the rotation. Second, *x*_*l*_ is added into the reconstruction for multiple (*N*_*s*_) times with parameters randomly picked from {*x*_*il*_}_1≤*i*≤*N*_. This is an inexplicit weighting strategy, called distribution weighting. The idea is that the contribution of *x*_*l*_ will be diluted when the support points has a wide distribution. *N*_*s*_ is typically set to 100 in current implementation.

### Rotation Using Unit Quaternion

The particle-filter algorithm needs to generate random support points and perform statistics for the distribution of the support points in the rotation subspace. This is difficult under the Euler angle system which is widely used in cryoEM. Therefore, we used unit quaternion to describe 3D rotation.

By definition, a rotation about the origin is a linear transformation of *ℝ* that preserves the origin, Euclidean distance and handedness. Once a basis of *ℝ* has been chosen, rotations can be represented by matrices. Especially, if orthonormal basis of *ℝ* is chosen, every rotation is described by an orthogonal 3 × 3 matrix with determinant 1. The quaternion is used to describe the rotation matrix.

Quaternion is an expansion of real number system, like complex. However, differing from complex number, quaternion has not one, but three imaginary parts, referred to as ***i***, ***j***, and ***k***. A quaternion has the form like ***q*** = *w* + ***i**x* + ***j**y* + ***k**z* = {*w, x, y, z*}.

A point *v* = {*x*_0_, *x*_1_, *x*_2_} can be described in quaternion system as *p* = {0, *x*_0_, *x*_1_, *x*_2_}. A rotation by a unit quaternion is 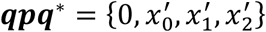 where ***q*** is a unit quaternion describing rotation with ‖***q***‖_2_ = 1, * indicates the conjugation and {*x*^*i*^, *x*^*i*^, *x*^*i*^ } is the rotated point. Moreover, assuming a rotation ***q***_1_ is followed by a rotation ***q***_2_, then the rotation will be 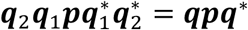, where the rotation quaternion is *q* = *q*_2_*q*_1_.

### Image processing

Three datasets were used for the tests. The dataset of cyclic-nucleotide-gated (CNG) channel is from our previous work^24^. The dataset of *Thermoplasma acidophilum* 20S proteasome^27^ (Entry code: 10025) and β-galactosidase in complex with a cell-permeant inhibitor^28^ (Entry code: 10061) were downloaded from EMPIAR^29^. All micrographs in movie mode are processed by MotionCorr2^11^ for motion correction and generating dose-weighted sum images. Relion1.4 was used for particle picking and extraction^14^. CTFFind3 was used for CTF determination^25^.

For the CNG dataset, the coordinates of 211,826 particles selected by 2D classification in previous work were used to extract the particles from 2820 micrographs with box size of 160 pixels, referred to as the dirty dataset. 87,149 particles processed by “polishing”^12^ of Relion1.4 from our previous work^24^, referred to as the clean dataset, were directly used for the further processing. The pixel size used in the image processing is 1.32 Å.

For the proteasome dataset and the β-galactosidase dataset, 112,412 and 89,857 particles are respectively selected from 131,319 and 108,226 picked particles by one round 2D classification, which is referred to as the dirty datasets. The coordinates of 49,954 proteasome particles and 41,104 β-galactosidase particles, downloaded from EMPIAR, were used to extract particles as the clean dataset. The initial pixel size and extracted particle box size are 0.6575 Å and 512 pixels for the proteasome and 0.637 Å and 768 pixels for the β-galactosidase, respectively. Finally, these particles were binned using the “Fourier cropping” method to 320-pixel box with 1.052 Å pixel size for the proteasome and 576-pixel box with 0.8493 Å pixel size for the β-galactosidase.

The 3D reconstructions were performed using the dataset generated above by a newly developed software implemented with the particle-filter algorithm. The final maps are sharpened with the post-processing method used in Relion^14^. The resolutions were measured based on the gold-standard FSC = 0.143 criteria.

**Supplementary Figure 1.**
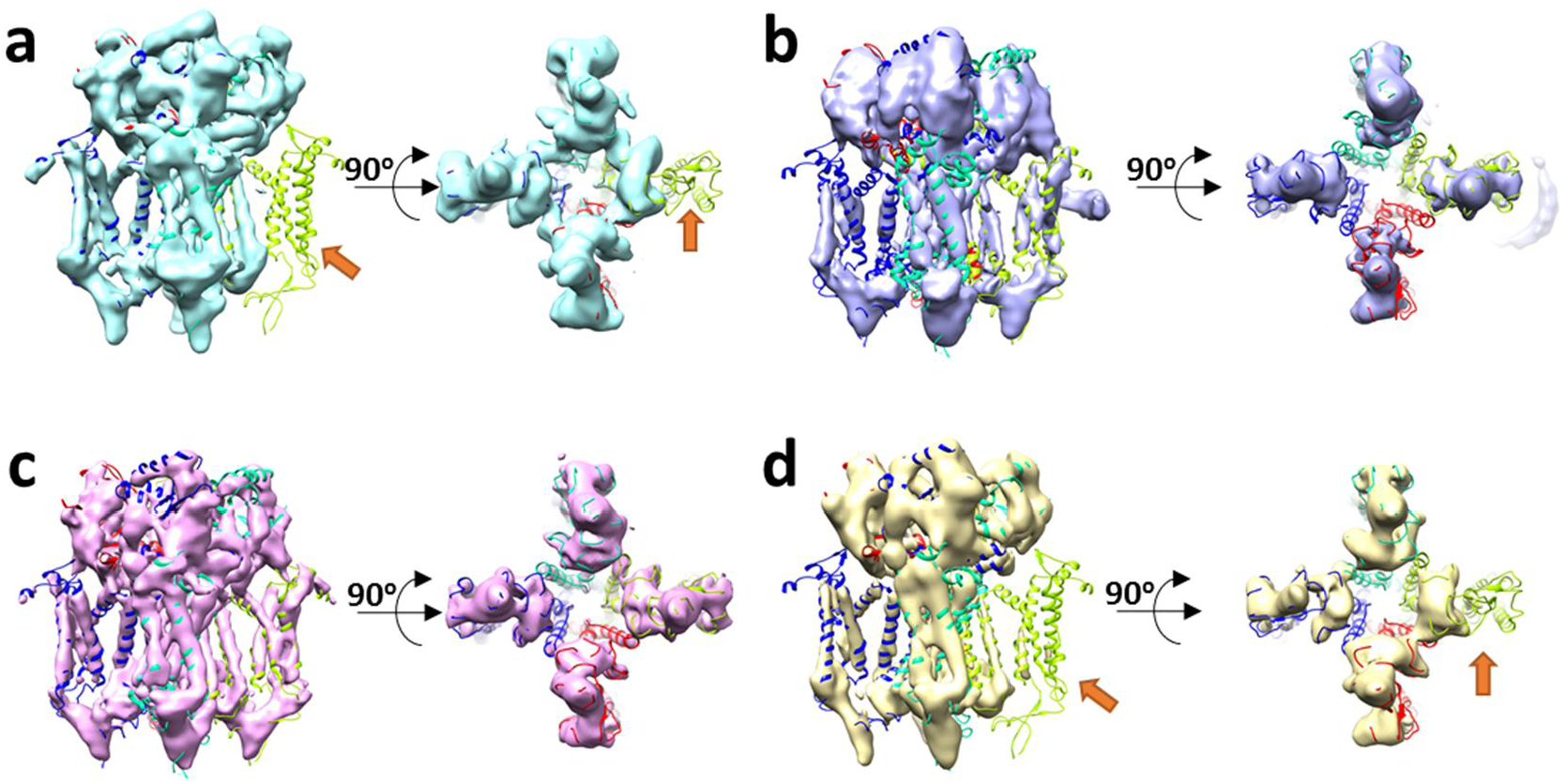
3D classification of the CNG dataset. **a**-**d** are four classes calculated from 211,826 particles after one round of filtering by 2D classification in our previous work^24^. The side view (left) and top view (right) are shown. The model with C4 symmetry (PDB entry code: 5h3o) was docked in the maps. The uncovered models, pointed by orange arrows, indicate the missing part of the subunit in the trans-membrane region of the CNG channel.

**Supplementary Figure 2.**
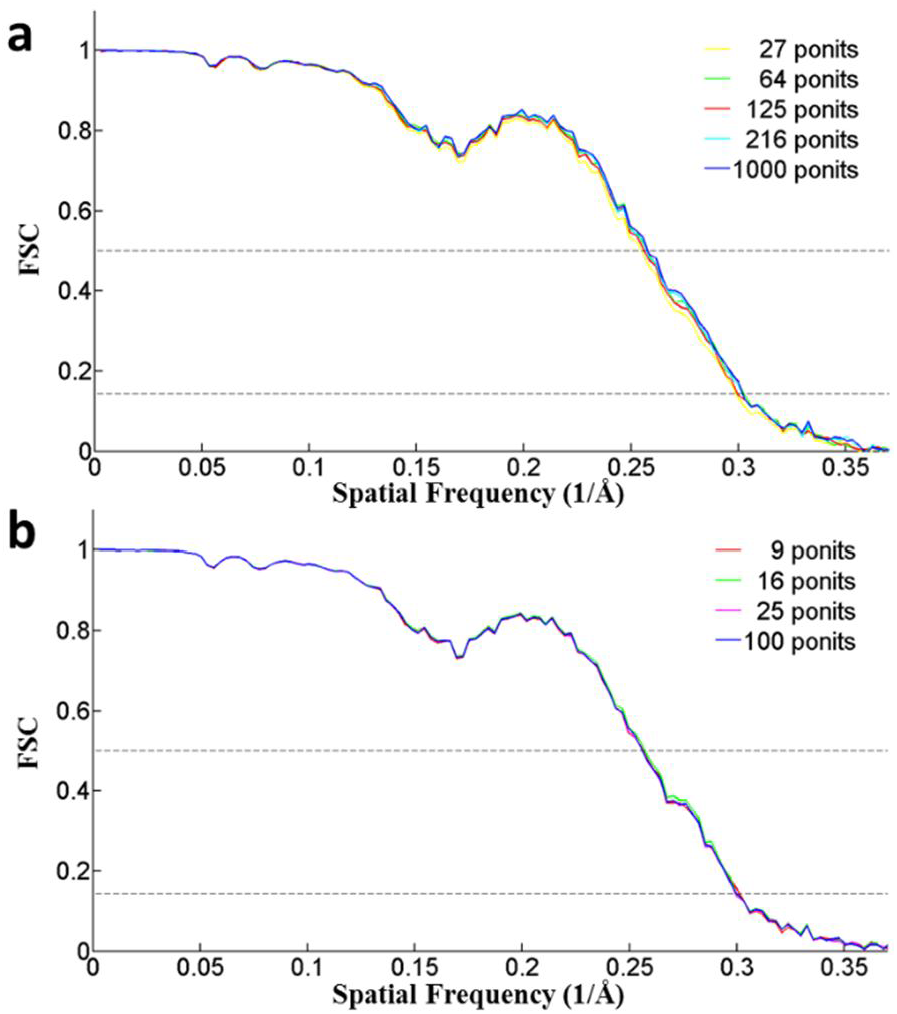
FSC curves of CNG reconstructions with various number of support points in **a**) the rotation subspace and **b**) x-y translation subspace. The nearly identical FSC curves with various number of support points implied that small amount of support points can give accurate parameter estimation.

**Supplementary Figure 3.**
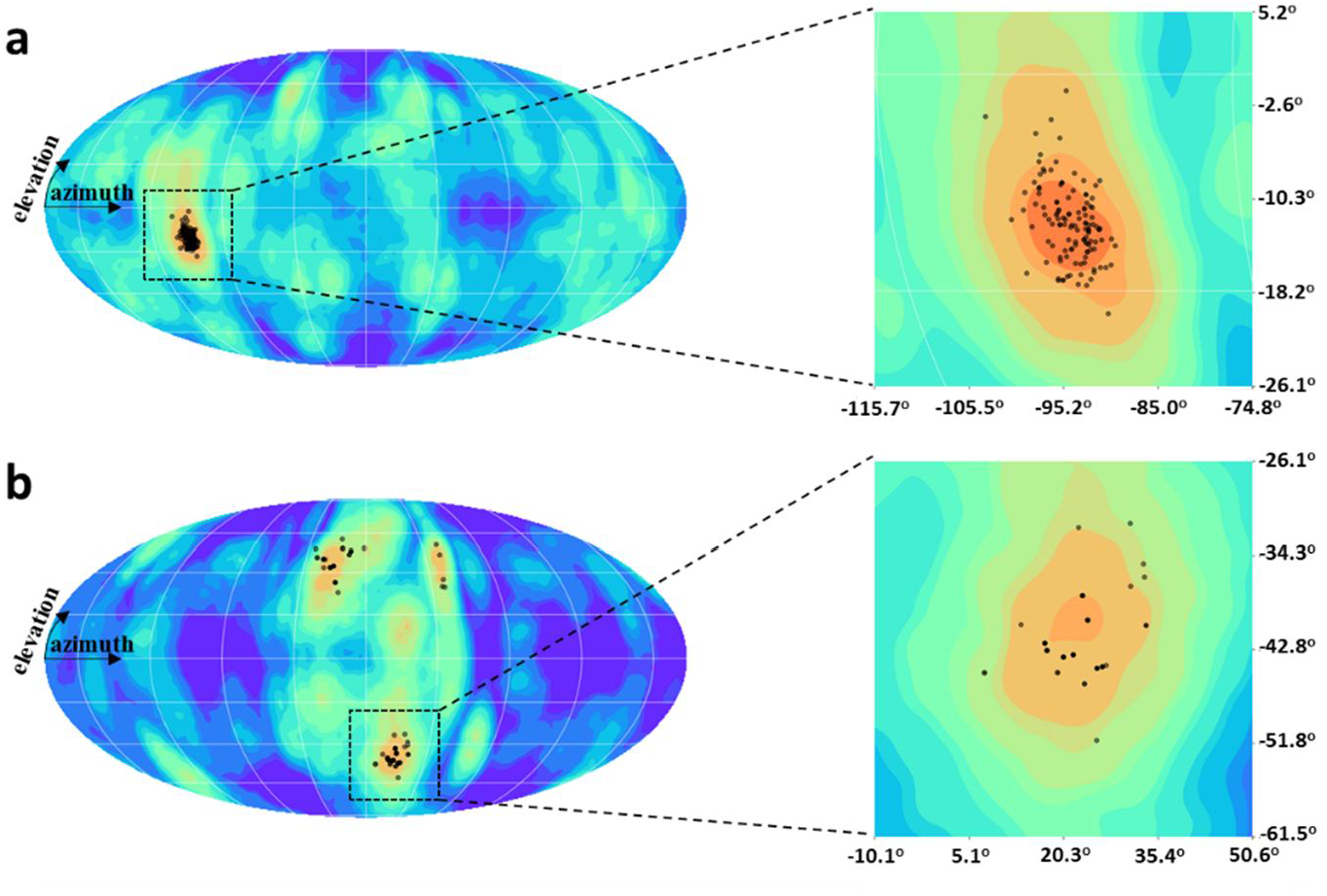
Support points on different likelihood functions. **a**) The support points distribute on a strong single peak of the LF which was calculated from a “good” particle image. **b**) The support points distribute on multiple strong peaks of the LF which was calculated form a “bad” particle images.

**Supplementary Figure 4.**
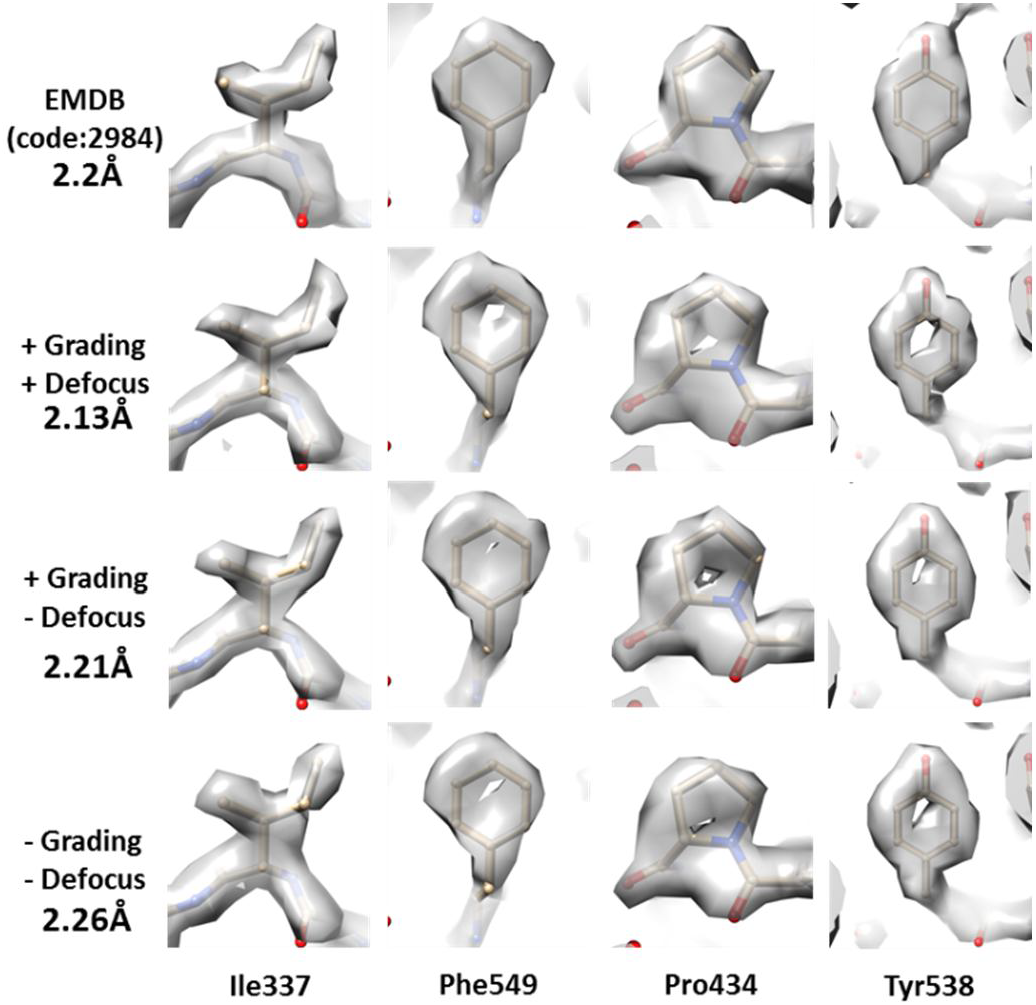
Representative side-chain maps of the β-galactosidase reconstruction using various options.

**Supplementary Figure 5.**
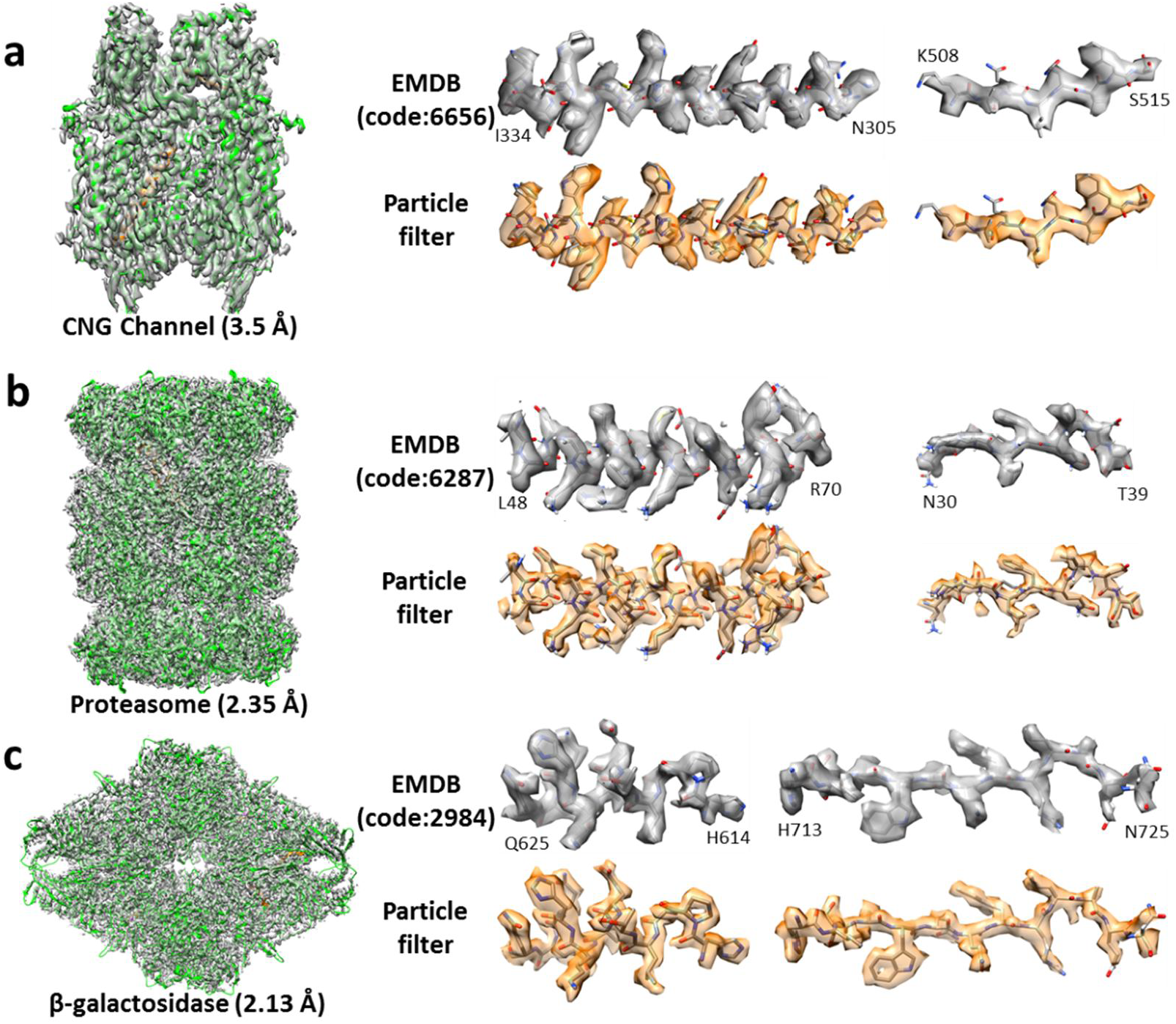
3D reconstruction of three datasets. **a**) the CNG density map, the proteasome density map and **c**) the β-galactosidase density map. The left images show the whole density maps with the resolution value labeled on the bottom, and the right are the representative secondary structures segmented from the maps deposited in EMDB and the maps on the left by the particle-filter algorithm.

## Appendix I

### Pseudo code of resampling algorithm

~~~
**Input**: 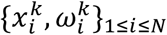
**Output**: 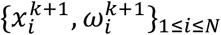
        Initialize the cumulative density function (CDF): *c*_1_ = 0
        **for** 2 ≤ i ≤ N **do**
                Construct CDF: 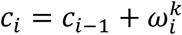
        **end for**
        Start at the bottom of the CDF: *i* = 1
        Draw a starting point from uniform distribution: u_1_~𝕌[0,1]
        **for** 1 ≤ j ≤ N **do**
                Move along the CDF: *u_j_* = *u*_1_ + (*j* - 1)
                **while** *u*_*j*_ *> c*_*i*_ **do**
                        *i* = *i* + 1
                **end while**
        **end for**
        Assign support point: 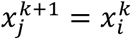
        Assign weight: 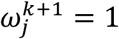
~~~

## Appendix II

### Pseudo code of the implemented particle-filter algorithm

~~~
**Input**: sampling space 𝕊, perturbation factor ε
**Output**: filtered support points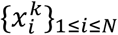, minimum support points standard deviation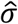
        **for** 1 ≤ i ≤ N **do**
                Initialize support points from uniform distribution: 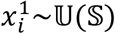
                Initialize weights: 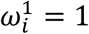
       **end for**
       **for** 1 ≤ i ≤ N **do**
        Initialize standard deviation: *σ* = ∞, 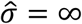
        **end for**
        Initialize measure index: *k* = 1
        **repeat**
                k = k + 1
                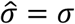
                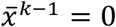
                **for** 1 ≤ *i* ≤ *N* **do**
                        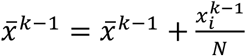
                **end for**
                σ = 0
                **for** 1 ≤ *i* ≤ *N* **do**
                        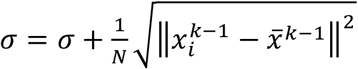
                **end for**
                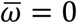
                **for** 1 ≤ *i* ≤ *N* **do**
                        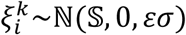
                        Perturb support points: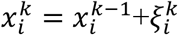
                        Calculate 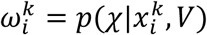
                        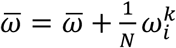
                **end for**
                **for** 1 ≤ *i* ≤ *N* **do**
                        Normalize: 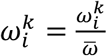
                **end for**
                Resample using algorithm in **Appendix I
       until** 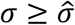
~~~

